# Measuring and modeling ecological rates with neutral theory

**DOI:** 10.64898/2025.12.02.691869

**Authors:** James G. Saulsbury, Scott L. Wing, Gene Hunt

## Abstract

The pace of change in ecological communities is central to classic and emerging problems in paleobiology and ecology. Yet measuring rates of change in community composition is problematic: when measured as ecological distance divided by time, rates appear faster when measured over shorter time intervals, preventing straightforward interpretation of rates when timescales vary. Similar problems arise when richness or sample size varies. We develop an approach that mitigates these issues using Hubbell’s neutral theory, a model in which species abundances change through stochastic births and deaths in a community with a fixed number of individuals *J.* Change in community composition is faster for smaller *J*, so 1/*J* provides a process-based rate metric that can be estimated from empirical time-series. We derive the likelihood for changes in community composition under neutral theory. Our approach accurately estimates rates in simulated data even when timescale, richness, and sample size vary. We use Quaternary pollen assemblage timeseries to demonstrate the power of this method for comparing rates between sites and through time and testing for rate shifts. Our approach can be flexibly adapted to test hypotheses on the causes of ecological change, including tests of coordinated stasis, neutral theory, and the timing of anthropogenic change.

## Introduction

The pace of change in community composition has a central place in both ecology and paleobiology. It features prominently in classical community ecological theory (Wangersky 1978), as well as in emerging problems involving anthropogenic ecological change, such as: is biodiversity declining (Johnson et al. 2024)? When did humans start to have a detectable impact on ecosystems (Mottl et al. 2021*a*; Nogué et al. 2021; Fastovich et al. 2025)? And what proximate causes mediate anthropogenic impacts on communities (Pinsky et al. 2025)? In paleobiology, ecological rates are likewise important: for example, the turnover-pulse/coordinated stasis theory predicts the existence of long-lasting blocks of community stability separated by brief intervals of rapid ecological and taxonomic turnover (Brett et al. 2025). Ecological rates also figure in the study of extinctions in the fossil record (Sheets et al. 2016; Smith et al. 2025), not least because extinction is the end result of population dynamics (Raup 1991; Liow and Stenseth 2007). Moreover, change in community composition has never been more broadly accessible to researchers through databases of ecological change like BioTime (Dornelas et al. 2025) or its paleontological equivalent, BioDeepTime (Smith et al. 2023).

Despite the conceptual importance of ecological rates and the availability of data for studying them, it is surprising how seldom they are actually computed. Although the literature commonly describes community change using rate-like terms such as slow, rapid, and abrupt, changes are almost always reported as magnitudes rather than as rates (magnitudes standardized by time) (e.g., Bobe et al. 2002; Wing et al. 2005; Birks 2007). The most intuitive metric of the rate of change in community composition, which we refer to non-pejoratively as a “naïve” rate, involves calculating ecological dissimilarity (for example, the Jaccard distance) between the composition of a community at two different times and dividing it by the relevant time interval (e.g., Bennett and Humphry 1995). Paleontologists have likely avoided computing naïve ecological rates because the field has long been aware that rate metrics for various processes can depend on the duration over which they are measured (Sadler 1981; Gingerich 1983). In virtually any real ecosystem, measured compositional change will appear slower the longer the timescale of measurement is. This is because community change is seldom purely directional, and when directional trends are reversed—say, when a rare species becomes abundant and then rare again—rapid short-term changes can cancel out to yield slow long-term changes (Collins et al. 2000). The resulting dependence of measured ecological rates on timescale creates serious problems for analysis by masking genuine biological rate variation. The problem is essentially model misspecification: comparing naïve rates calculated on different timescales implicitly assumes a model where change scales linearly with time, but reality is not like this. As a result, the only paleontological systems for which ecological rates are commonly measured are those with dense temporal coverage that allow users to interpolate or bin samples into intervals of constant temporal duration. Most such studies analyze Quaternary pollen assemblage timeseries, where rate calculations are often combined with a smoothing step to reduce noise (rate-of-change analysis; Jacobson and Grimm 1986, Mottl et al. 2021*a* and references therein; see also Gibbs et al. 2012).

As in paleoecology, studies of contemporary ecological change compute rates only rarely. Approaches in this area are diverse (e.g., Pinsky et al. 2025; Nwankwo and Rossberg 2026), but perhaps the most common involves measuring temporal variability (Gaston and McArdle 1994; Cottingham et al. 2001; Dornelas et al. 2014; Montgomery et al. 2026). Like naïve rates, measurements of temporal variability cannot be straightforwardly compared when timescale varies, and for similar reasons: comparing temporal variabilities across different timescales relies on the unrealistic assumption that change is independent of timescale (Pimm and Redfearn 1988). Across contemporary and paleontological studies of ecological change, additional headaches arise from the effects of taxonomic resolution, rare species (Overpeck et al. 1985), and incomplete sampling (Link and Nichols 1994) on measures of change. In short, there remains no general solution to the problem of comparing rates of ecological change under variable sampling conditions, making it challenging to find statistical support for claims about the pace of ecological change.

In this paper we develop an approach to measuring rates of change in community composition based on a model of neutral ecological drift and demonstrate our approach with data from Quaternary pollen assemblages. Our goal is to be able to estimate the rate of ecological drift that best explains an observed change or set of changes in community composition. This will solve the rate-time scaling problem to the degree that neutral theory provides a realistic picture of how rate scales with time, and as a process-based model it can simultaneously solve other problems such as incomplete sampling. Several studies have used simulations of neutral ecological change for studying empirical temporal turnover (McGill et al. 2005; Reymond et al. 2011; Sgardeli et al. 2016; Lewandowska et al. 2020) and for studying general relationships between turnover patterns and neutral parameters (Nakadai 2025), but to our knowledge ours is the first to approach the problem analytically. Our method mitigates the major problems in measuring ecological rates and creates a likelihood-based framework for testing hypotheses on the causes of ecological change in modern and fossil communities.

### The Model: Ecological Drift

We model community change as the local ecological drift component of Hubbell’s (2001) unified neutral theory of biodiversity. Hubbell’s neutral theory is a quantitative representation of the dynamics of a community with a fixed number of individuals *J* belonging to different species having complete niche overlap and equal fitness. At each unit of time (say, each year), a new set of species abundances are drawn as a multinomial sample of size *J* defined by the species proportions in the previous time step. Because this is a random sample, the proportions will differ slightly from year to year and community composition will drift over time. Once a species has zero abundance, it does not occur again in the community. Our approach differs from Hubbell’s full model in ignoring the possibility of speciation and migration from a metacommunity, and also in how individual deaths are scaled with respect to time. In Hubbell’s version, each time step represents the death of single individual (a variant of the Moran process; Etienne and Alonso 2007), but in the present treatment, tractability was achieved by assuming all individuals die at the same time and are replaced all at once (the Wright-Fisher process). The Moran and Wright-Fisher processes converge on the same predictions for abundance change when *J* is large (Blythe and McKane 2007; Etienne and Alonso 2007), so our model is equally applicable to organisms with overlapping or non-overlapping generations. The model only applies to relative abundances (proportions of individuals in a community belonging to each taxon) and not absolute abundances (e.g., census counts). This is because we want a metric that can be compared across the timescales of modern and fossil change, and while the absolute abundances of fossil organisms are generally not knowable (though see Marshall et al. 2021), relative abundances are often faithfully preserved in fossil assemblages (Tyler and Kowalewski 2023).

The single parameter of this model—total community size, *J*—determines the pace of compositional change in the community. This is because compositional change in this model is fundamentally a sampling process, and like any sampling process, the amount of noise decreases with sample size. In a familiar example, the average proportion of heads in 1,000 coin tosses will be very close to 0.5, but this proportion will be much more variable when only 10 coins are tossed. Likewise, when community size *J* is high, species proportions in one generation will differ only slightly from those of the previous generation. At low values of *J*, past relative abundances will be less faithfully represented in the next generation, resulting in larger changes from one time step to the next. Cumulated over time, the relative proportions of species will change exactly as do haploid allele frequencies experiencing neutral genetic drift. Higher rates of community change at lower *J* are analogous to the greater potency of genetic drift in small populations compared to larger ones. By analogy with the way rates of drift are dealt with in population genetics (Hu et al. 2006), in the following we consider drift rate as 1/*J*.

The parameters of Hubbell’s model are usually inferred from the static relative abundance distribution of species (Hubbell 2001; McGill 2003; Volkov et al. 2003; Etienne and Olff 2004; McKane et al. 2004; Olszewski and Erwin 2004; but see Gotelli and McGill 2006). However, because we are interested in the pace of community change over time, we wish to fit this model using temporal changes in species abundances. One approach involves comparing observed patterns with simulations (McGill et al. 2005), but this is often prohibitively slow and requires information, such as total community size, which is unconstrained in most empirical studies. The approach we develop here allows for fast analytical estimation of *J* using only a record of changes in relative abundances for species in a community. The approach is based on multinomial sampling theory, but with four extra steps to make the problem analytically tractable: (1) change in a community of size *J* over *t* generations is estimated as change in a community of size *J*/*t* over one generation; (2) the continuous binomial distribution is used instead of the standard binomial distribution; (3) changes in multiple species are described by combining multiple binomial distributions using conditional probability; and (4) incomplete sampling is accounted for by modifying *J/t*.

### Model Implementation

#### 1. Substituting a single generation for many

Under neutral drift in a local community with non-overlapping generations, the abundance of a species at time *t* = 1 is given by the binomial distribution with size parameter *J* and probability equal to relative abundance of species at time *t* = 0. Considering change over multiple generations is more complicated. Hubbell (2001) analyzes abundance changes over multiple generations by building a transition matrix and taking it to a power *t*, but the difficulties of multiplying large matrices together means this not feasible for *J* larger than a few thousand. It also rules out jointly considering changes in multiple species. (We do use transition matrices to ground-truth our own approach; Fig. 1A, black lines.) Our solution starts with the recognition that, under a range of circumstances, the rate of neutral drift scales with the ratio of community size to time. More specifically, Azaele et al. (2016, Eqn. 32) show that, when relative abundances are not too close to 1 or 0, are recast as continuous, and change via diffusion, their rate of drift depends on *J/t*, regardless of *J* or *t* individually. We take advantage of this unique feature of drift to use a community of size *J/t* changing over a single generation as a kind of scale model of a community of *J* individuals changing over *t* generations. The probability that the relative abundance of a species changes from *x* to *y* after *t* generations can therefore be approximated using a binomial distribution with size *J/t*:

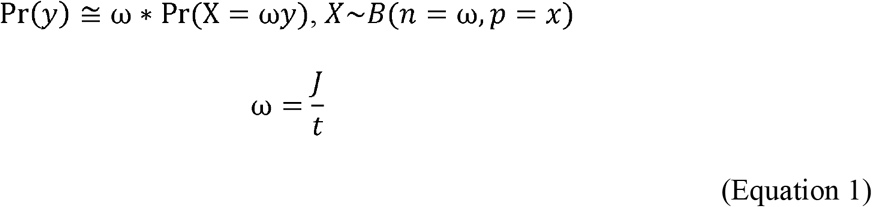

where *B(n,p)* is the binomial distribution with size parameter *n* and success probability *p*. We multiply the resulting probability by ω to rescale the probability density function to have support for *y* on the interval (0,1), since our interest is in relative abundances. Equation 1 provides excellent approximations of true change probabilities as estimated with transition matrices (Fig 1, red dots). However, it does not accurately capture true probabilities of extirpation, especially as extirpation becomes probable; our approach therefore ignores species transitions that end in zero abundance (see next section). Because the local drift model does not include speciation or migration, we also ignore species transitions beginning at zero abundance.

**Fig. 1.**
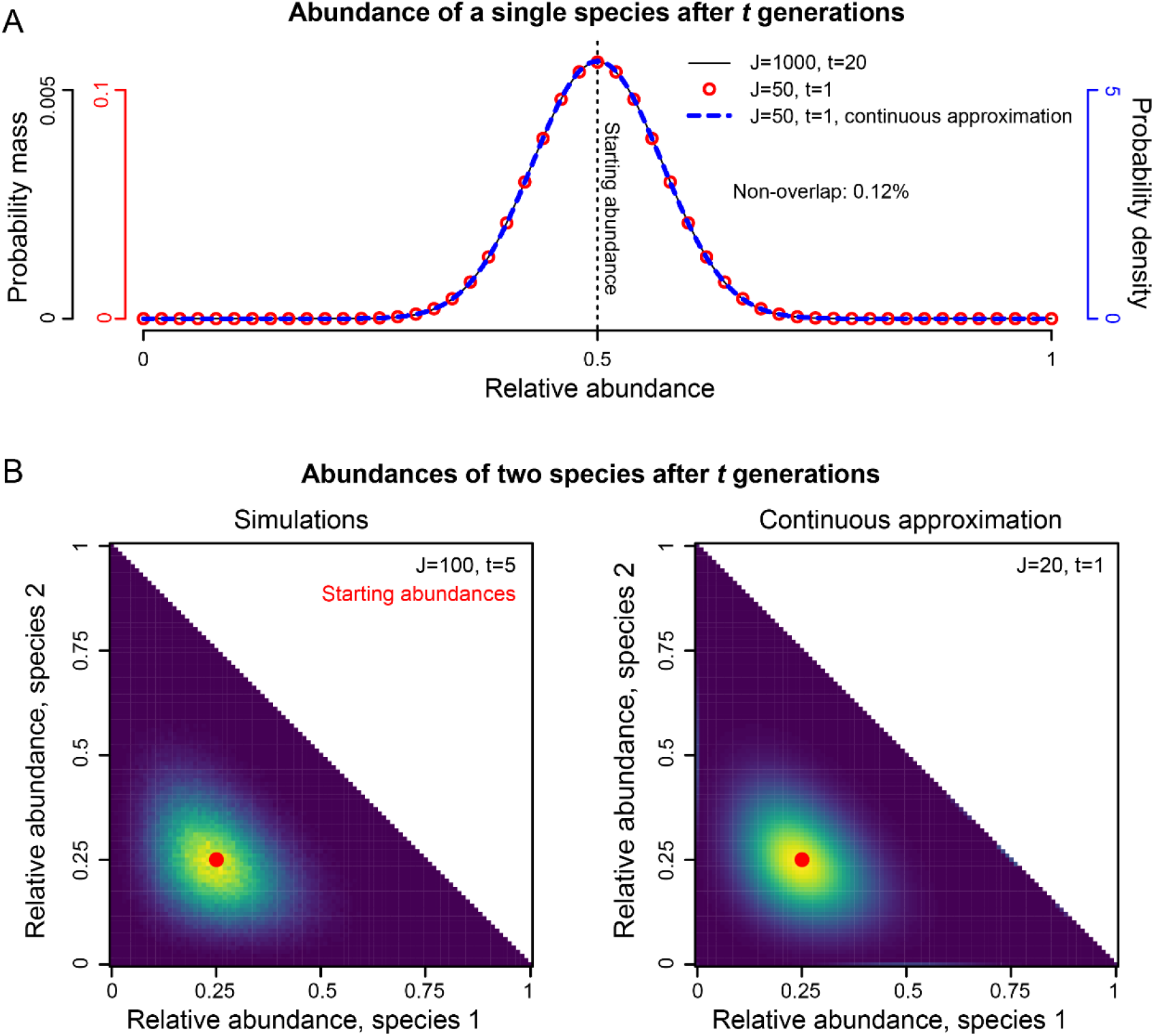
Estimating the likelihood of observed changes in community composition under neutral theory. **1A.** Probability of a species having a given relative abundance after t generations of ecological drift. The expected change in a large community over many generations (solid line) is nearly identical to expected change in a small community over one generation with the same ratio of J (community size) to t, expressed either with the binomial distribution (red dots) or with a continuous binomial distribution (blue dashed line). Also shown is percent non-overlap between true probabilities and continuous approximation. **1B**. Probability of two species having any set of relative abundances after t generations, estimated through simulations (left) and through Eqn. 3 (right). Brighter colors represent higher probability.

#### 2. Substituting continuous for discrete binomial distribution

Eqn. 1 is not defined when ω (the size parameter in the binomial distribution) and ω*y* (the number of successes) are not whole numbers. This makes most empirical relative abundances and most values of *J* impossible to consider. To get around this, we replace the binomial distribution with the continuous binomial distribution, which spreads the probability density at each value *i* across the interval [*i*,*i*+1] (Ilienko 2013). The cumulative distribution function (CDF) of the discrete binomial at *i* is equal to CDF of the continuous binomial at *i*+1, and the probability mass of the discrete binomial at *i* very closely matches the probability density of the continuous binomial at *i*+0.5. Thus,

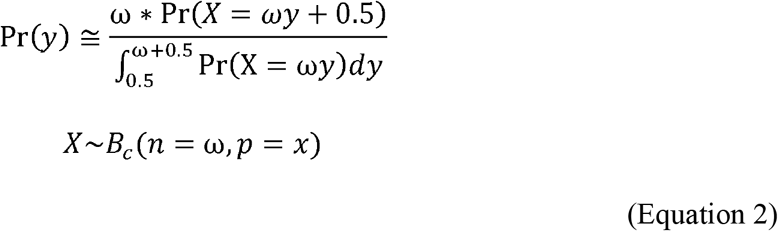

where *B_c_*(*n*,*p*) is the continuous binomial distribution. Because *X* ranges from 0 and size + 1 but transformed relative abundance ωy + 0.5 can only take values between 0.5 and size + 0.5, we divide by the integral in the denominator so that probabilities from this expression sum to 1. We also found that the complement of this integral corresponds closely, though in a biased fashion, to true extirpation risk (slope = 0.80, R^2^ = 0.994; Fig. S3). Eqn. 2 therefore approximates the probability of abundance change conditional on a species not going extinct, which is important since our method ignores local extirpations. In general, Eqn. 2 closely matches true probabilities of change (e.g., Fig. 1A shows only 0.12% non-overlap between true and approximated probabilities).

#### 3. Multiple species

The joint probability of a set of abundance changes in multiple species differs from the product of probabilities calculated independently for each species because changes in relative abundance in different species are negatively correlated with each other. Since our scaled-down community of size *J/t* changes by sampling with replacement over a single generation, we can calculate the joint probability of multiple abundance changes with the multinomial distribution. We express this as the product of multiple binomial distributions modified using conditional probability (SI).

Thus, the probability of a set of relative abundances Y for *S* species at time *t* is given by:

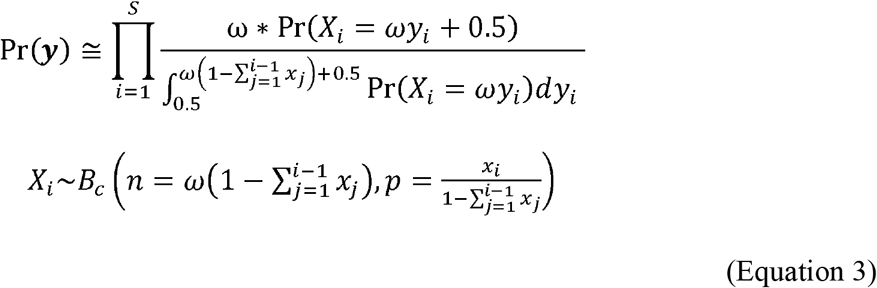

where *B_c_* is the continuous binomial distribution and *x_i_* and *y_i_* are the relative abundances of species *i* at times 0 and *t*, respectively. The abundance change in each species *i* is considered for a community in which species 1 through *i*-1 have already been considered and removed, reflected in the sigma terms in Eqn. 3. If the sets of relative abundances x and Y both sum to 1, the change in the last species is fully determined by the changes in the other species, in which case we omit the last species from calculation. The likelihood of a time-series is the product of Eqn. 3 calculated for every observed change in community composition. This approach closely approximates the results of simulations, including capturing the negative covariation between changes in multiple species (Fig. 1B). Because our approach calculates the probability of the model for each consecutive change in community composition, our method can be naturally extended to estimate the rate of change not only for an entire timeseries but also for each step in that timeseries.

#### 4. Incomplete sampling

Observed relative abundances at times 0 and *t* can differ from each other due to two kinds of processes: first, the genuine biological change in composition between times 0 and *t*, and second, the sampling processes that add noise to our estimates of true species abundances at times 0 and *t*. We assume the observation process happens via sampling with replacement. This is realistic for cases in which a single biological individual can leave multiple occurrences, including pollen and most plant body fossils, trilobites and other molting animals, and vertebrate teeth. We suggest our approach is equally applicable to other groups because the results of sampling with and without replacement only diverge when the number of samples approaches the number of individuals, which is rare in ecological datasets and perhaps never occurs in paleoecology. Because true change in the neutral model also happens by sampling with replacement, each observation step can be treated as if it were an additional phase of drift with *J/t* value equal to the sample sizes. These three steps of drift can be combined into a single step (SI), resulting in a multinomial distribution with size equal to the reciprocal of the sum of reciprocals for the three steps:

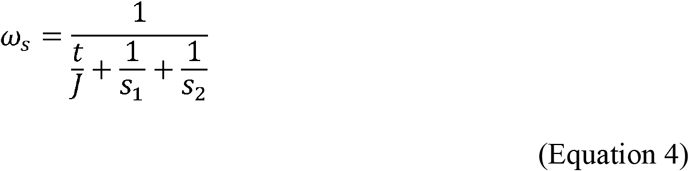

where *s_1_* and *s_2_* are sample sizes for the communities at times 1 and 2. w_s_ can be substituted for ω in Eqn. 3 to model incomplete sampling. Note that, as sample sizes get very large, w_s_ converges on *J/t*, so with perfect sampling the problem reduces to modeling true change.

#### Software

We implemented our approach in a new R package, *ecoDrift (*available at https://github.com/jgsaulsbury/ecoDrift). The main function *fitJ*() uses the *optimize*() function in the ‘stats’ package (R Core Team 2026) to find the value of *J* that maximizes likelihood for an observed time-series. Fitting *J* to an empirical time-series requires an assumption about generation time in the system being studied; the default is 1 year. *fitJ*() can also provide a profile confidence interval: it uses a chi-square table to find the range of *J* corresponding to a given α, returning a 95% confidence interval by default. *fitJ*() is fast, taking about 3s to run on a personal computer with a time-series of 100 species and sampled at 100 different times. Runtime is proportional to the number of species minus one and the number of sampled time intervals minus one (Fig. S4). We also provide the function *plot_Js*() for the purpose of visualizing how ecological rates change through time. This function fits *J* independently to each transition, and plots 1/*J* values because these are more easily interpreted as rates. The *simDrift*() function simulates neutral community change, with the possibility of including migration from a static metacommunity. Finally, the *fitJshift*() function fits a 3-parameter model to a dataset in which *J* undergoes a shift at some point in the time-series.

### Method Performance: Likelihood Approximation and Parameter Estimation

We simulated neutral change in an even community of size J = 10^5^ under a variety of timescales, sampling intensities, and species richness values, and tested the ability of our approach to recover true *J*. For comparison, we also measured naïve rates—ecological distance divided by time—for the same simulations. We measured Jaccard distance (one minus intersection over union) because it is widely used and intuitive, but in general our findings apply across distance metrics. First, we simulated drift in a perfectly sampled community with 20 species over an interval ranging from 1 to 200 generations. Naïve rates showed strong negative scaling with timescale: average log rates decreased by roughly 14 standard deviations over an order-of-magnitude increase in timescale (Fig. 2A). Conversely, drift rate 1/*J* was estimated accurately regardless of timescale (Fig. 2D). Next, we evaluated the effect of sample size by simulating drift over 100 generations in a 20-species community and drawing N samples with replacement from the community before and after the change, with N varying from 100 to infinity (perfect sampling). Lower sample size results in higher naïve rates as community change comes to be dominated by incomplete sampling (Fig. 2B). No such effect is seen in estimated drift rates: median estimated 1/*J* matches the true value across all sampling conditions (Fig. 2E). However, when sampling is low (in this case, below N = 500), observed community change can sometimes be entirely explained by sampling noise, which can drive estimates of drift rate (1/*J*) to 0. Lastly, we simulated drift in communities with initial species richness varying from 2 to 500. We supposed that more diverse communities would exhibit faster apparent rates of change because more abundance contrasts can be observed in communities with more taxa (Bennett and Humphry 1995). Results confirmed this suspicion: naïve rates are higher in more diverse communities, even though the generating drift rate was the same in all cases (Fig. 2C). Average estimates of 1/*J* do not scale with number of taxa (Fig. 2F). Precision in estimating 1/*J* increased with both sample size and number of taxa.

**Fig. 2.**
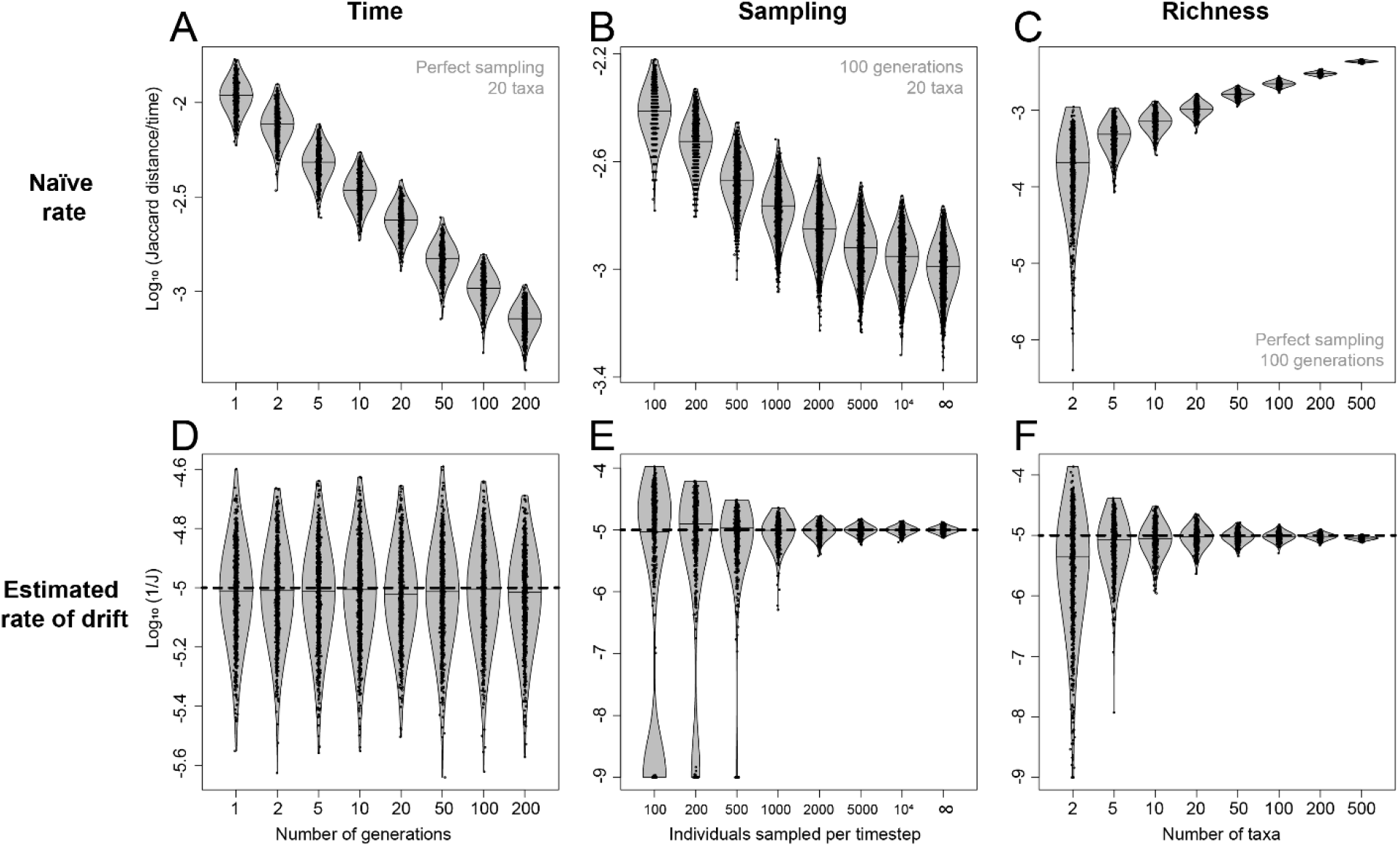
Naïve rates (**2A–C**) and estimated drift rates (**2D–F**) fit to simulated data under differing amounts of time (**2A, D**), sampling intensity (**2B, E**), and richness (**2C, F**). Naïve ecological rates scale with time, sampling intensity, and richness, while estimated drift rates do not. Dashed line indicates true rate in bottom row. Each violin visualizes rates from 10,000 simulations using normal optimal smoothing kernel density; lines are medians.

We also tested the range of true drift rates that could be accurately estimated using our method. We did this by simulating time-series under different values of 1/*J* for a total length of 2000 generations, sampled every 100 generations with sample sizes of 2000. 1/*J* could be accurately estimated for true 1/*J* spanning over three orders of magnitude (Fig. 3). At very low rates (high *J)*, observed community change was dominated by sampling noise (Fig. 3, left inset), causing information on true rate to be lost. At very high rates (low *J*), intense extirpation likewise results in loss of information and wide variance in rate estimates.

**Fig. 3.**
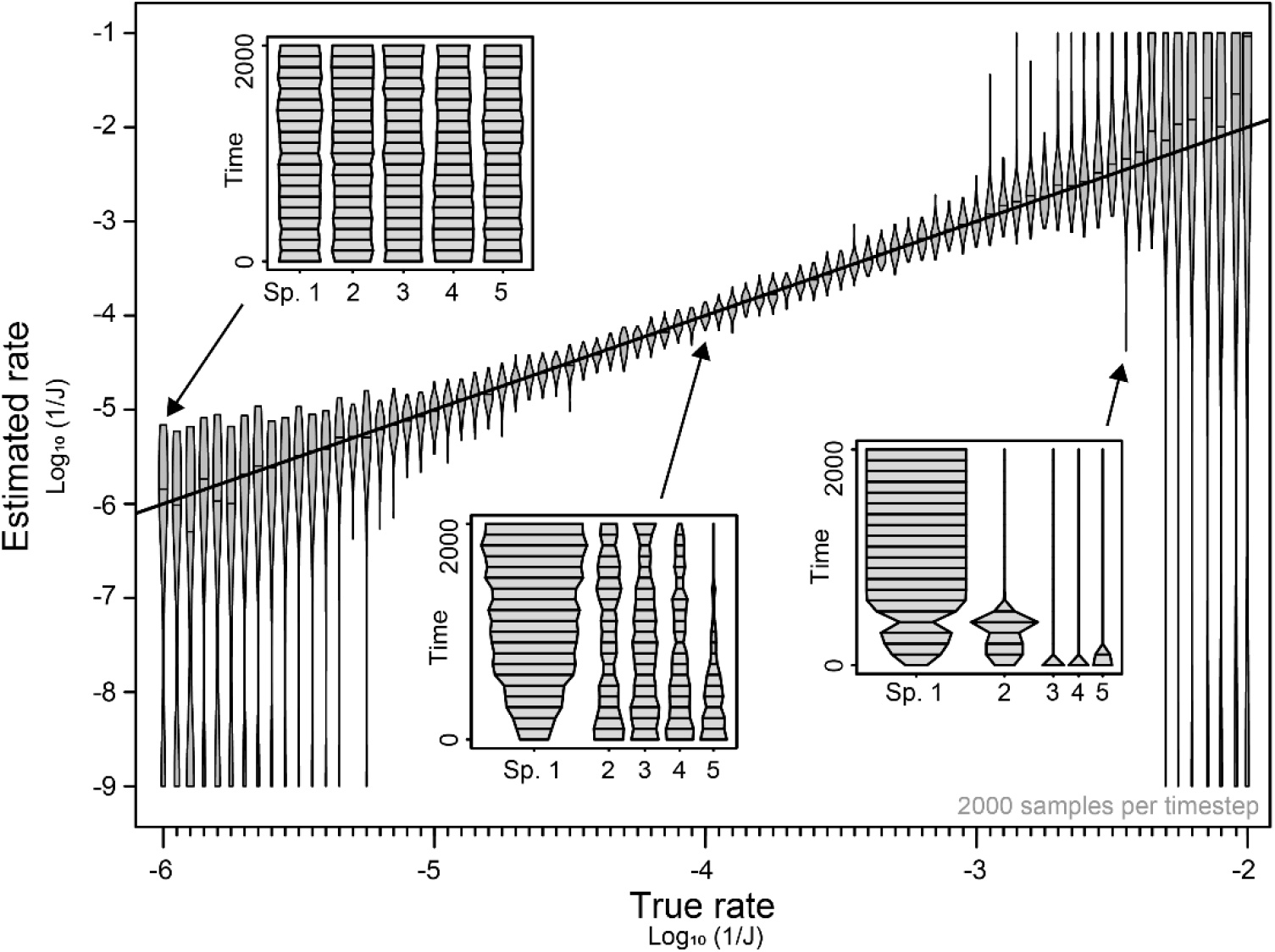
Violin plots of true vs. estimated rates of drift across four orders of magnitude of 1/J values, along with a one-to-one line. Insets show examples of time-series simulated with low, medium, and high rates. Violins as in Fig. 2.

Finally, we systematically explored the ability of our method to capture true probability distributions and accurately estimate drift rates across the range of conditions encountered in real community composition data. We took records of Quaternary pollen assemblage composition from the Neotoma database (see next section) as representative of paleoecological timeseries. We considered performance across the axes of relative abundance and drift amount, obtained by fitting drift rate 1/*J* to all >70,000 community transitions in the dataset and multiplying by *t* to consider the amount rather than the rate of drift. We chose computationally convenient bounds on these axes encompassing roughly the middle 95% of both distributions: thus, we visualized the representation in the Neotoma database of relative abundances ranging from 0.001 to 0.63 (the 2.5^th^ through 96^th^ percentiles) and of drift amounts ranging from *t/J* = 0.0005 to 0.1 (the 2^nd^ through 97.6^th^ percentiles) (Fig. S2A). We used a transition matrix with *J* = 2000 to calculate true abundance probability distributions across this parameter space, then compared these with probability distributions estimated by our method using percent non-overlap. We also simulated data across parameter space in a two-species community of size *J* = 20,000, fit 1/*J* to 2,000 such simulations for each set of conditions, and measured error as log_10_(estimated 1/*J*)-log_10_(true 1/*J*).

Error in estimated probability distributions (Fig. S2C) and drift rates (Fig. S2D) scale roughly with the probability of extirpation. When abundance is low enough and drift high enough for extirpation risk to exceed 75%, non-overlap between true and estimated probability distributions exceeds 10% and 1/*J* is overestimated by 0.15 to 0.44 orders of magnitude. However, outside this small corner of parameter space, both forms of error are minimal. Average non-overlap, weighted by frequency of empirical observations in each part of parameter space, is 4.1%, and the weighted average of absolute error in estimated 1/*J* is 0.062 orders of magnitude. Our method therefore performs well across a breadth of realistic conditions, and we expect it can be useful for extracting information on ecological change for a variety of taxonomic groups, sampling setups, and spatiotemporal scales.

### Empirical example: Quaternary pollen

#### Data

We demonstrate the utility of our approach by fitting ecological drift models to Quaternary pollen assemblage data from the Neotoma database (Williams et al. 2018), which has long been used to study floral change over the last ∼100,000 years. The pollen assemblages record changes in plant community composition over annual to thousand-year timescales, although the relationship between pollen abundance and local plant abundance is not one-to-one (Bradshaw and Webb III 1985). We selected pollen datasets from two cores with comparable temporal resolutions but with different apparent ecological dynamics to demonstrate the use of our model for identifying differences between time-series. We selected a third with an apparent increase in ecological rate near the top of section to demonstrate the use of our approach to test for a rate shift. Finally, to examine rate-time scaling across the Neotoma database, we accessed the 1,362 sites from Fastovich et al. (2025), which we subset further to the 1,250 sites for which those authors were able to harmonize taxonomies and standardize age models. In all exercises, we modeled communities as if all species had a generation time of one year. Investigators may choose to use a different generation time that more closely reflects their study system, but we suggest that maintaining annual turnover as a part of the idealized neutral community will allow for more straightforward rate comparisons between systems, especially given that estimates of generation time will often be variable and imprecise (as in evolutionary studies; Hansen 2024).

#### Comparing time-series

We plotted relative abundance through time for pollen taxa from two time-series (Fig. 4). The first, Secret Valley Marsh in northeastern California, USA, encompasses 38 sampled horizons across 9,926 years, with an average temporal resolution of 269 years and average sample size of 744 specimens; the site is an upland chaparral dominated by grass and sedge pollen and with a history of burning. The second, Vuolep Allakasjaure in northern Sweden, is an alpine site above the treeline with 80 sampled horizons, an average temporal resolution of 243 years, and an average sample size of 591 specimens; birch and pine pollen are common. Secret Valley Marsh appears to have more volatile community dynamics, with numerous shifts in rank abundance order. Vuolep Allakasjaure appears to change more slowly, with gradual changes upsection including a decline of birch pollen and an increase in pine and sedge pollen.

**Fig. 4.**
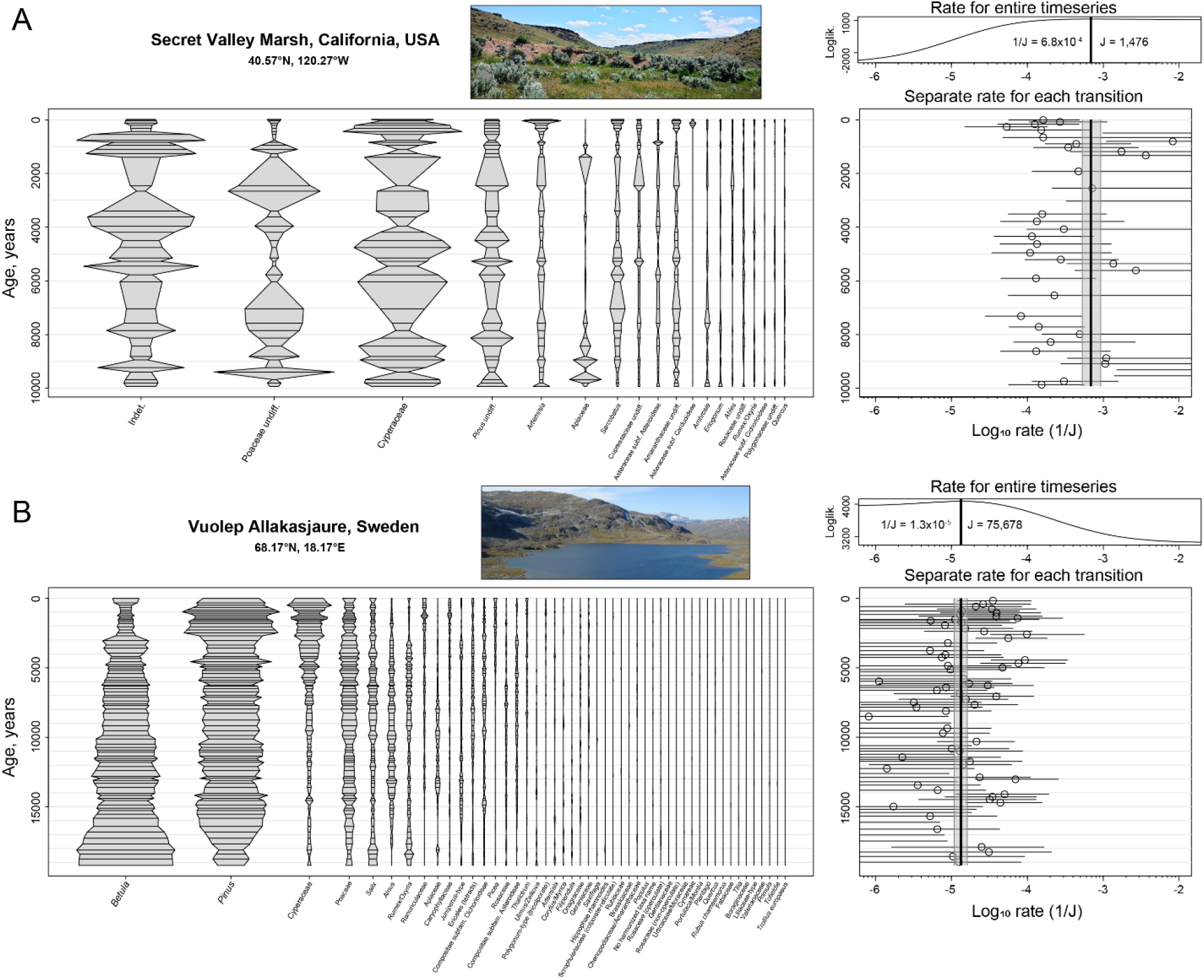
Fitting rates of drift to two pollen abundance time-series. For each time-series, we plot relative abundances of all taxa through time, the likelihood surface for a single rate of drift fit to the entire time-series, and the results of fitting rates separately for each transition. Error bars in bottom right panels show 95% confidence intervals for each transition. Grey envelope around bold vertical bar indicates 95% confidence interval for entire time-series. **4A.** Secret Valley Marsh. Image © W. Johnson. **4B.** Vuolep Allakasjaure. Image from Jonsson (2009).

Fitting drift models to these two sites confirms the contrast between them: best-fit 1/*J* values for the Secret Valley Marsh and Vuolep Allakasjaure sites are 6.8×10^-4^ and 1.3×10^-5^ respectively, corresponding to a ∼50-fold difference in rates (Fig. 4). Fitting 1/*J* separately for each transition reveals temporally autocorrelated rate variation across both time-series, but especially at Secret Valley Marsh. Rate does not correspond closely to temporal resolution: for example, high temporal resolution in the past 1000 years corresponds to low rates of change at Secret Valley Marsh. Uncertainties around 1/*J* estimates for the Vuolep Allakasjaure dataset are broader especially in the direction of lower rates, likely because of the phenomenon highlighted in Fig. 2E where very low rates can combine with incomplete sampling to yield a change that is indistinguishable from stasis.

#### Testing for a rate shift

We next tested for a shift in the rate of change at the Saint-Thomas site near Nante, France (Fig. 5). Oak and alder pollen dominate throughout the time-series, but in the last two millennia several other taxa became common including sedges and grasses (Fig. 5A). Fitting 1/*J* independently to each transition (Fig. 5B) reveals many high-rate transitions in the last two millennia, and many intervals with no detectable change before that time. We fit a rate shift model to this time-series with three parameters for a first and a second rate as well as for the age at which this transition occurs. We find the time-series is best described by a transition 1,717 years before present from a low rate of change (*J* = 6.7×10^-5^) to a rate almost 9 times faster (*J* = 5.8×10^-4^) (Fig. 5B, bold line). As in the previous example, estimated rates do not correspond to temporal resolution: the highest resolutions are between seven and five thousand years ago, when estimated drift rates are relatively low. Fig. 5C shows likelihood profiles for the age of the rate shift, obtained by finding best-fit values for the first and second drift rates for each possible shift age. The best model with a rate shift at 1,717 years before present improves on a one-rate model with ΔAIC = 243. Thus, model-fitting can yield decisive tests of hypotheses on rate changes in the presence of timescale variation.

**Fig. 5.**
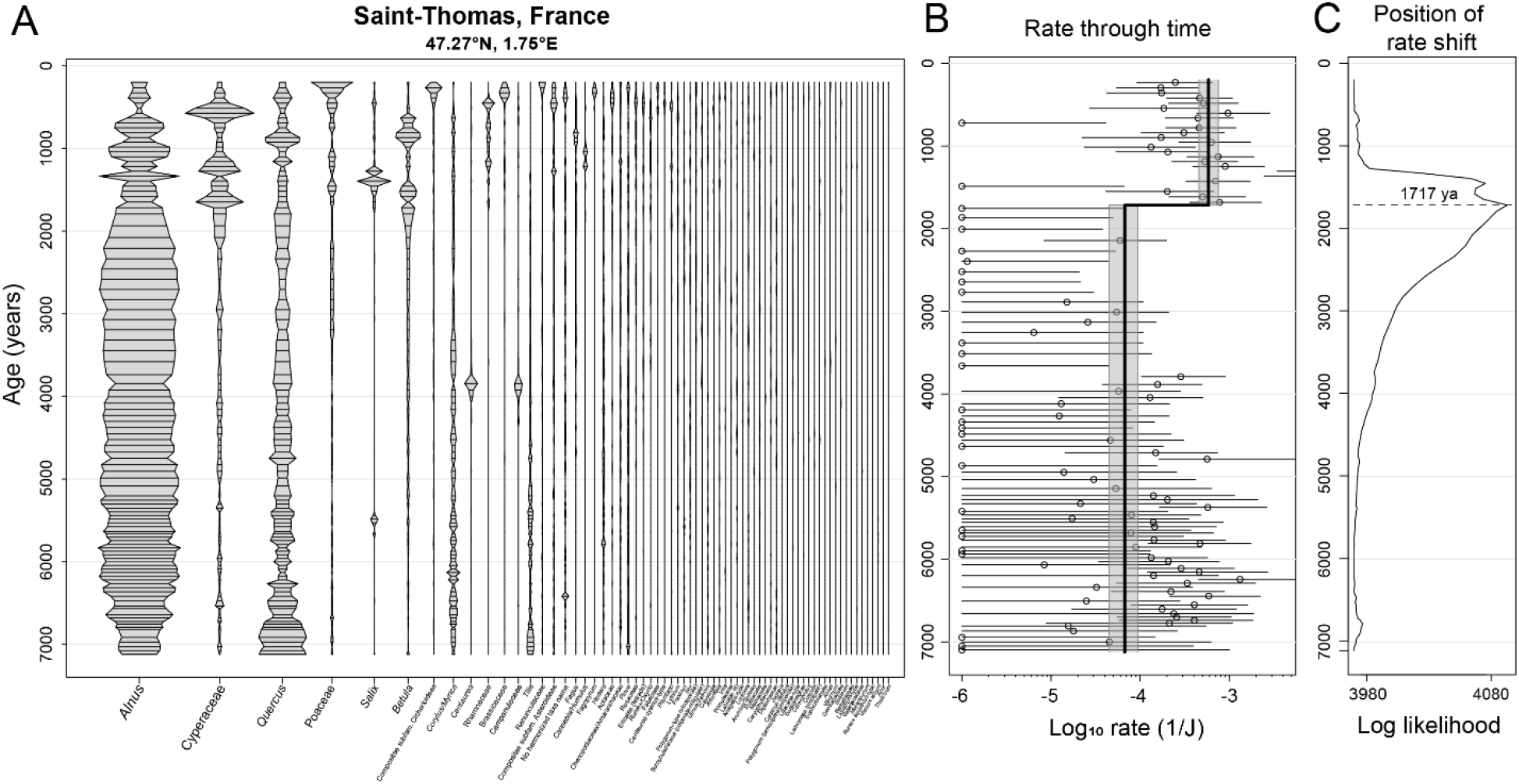
Investigating a time-series with an apparent increase in the pace of change. **5A.** Relative abundance of all taxa through time at Saint-Thomas, France. **5B.** Rate of drift through time. Points and error bars show rates fit separately to each transition as in Fig. 4. Bold line shows the best-fit two-rate model. Grey envelope shows simultaneous 95% confidence interval on the two rates and the time of transition. No other transition times are within the confidence interval. **5C.** Likelihood surface for the position of the rate shift.

#### Empirical timescaling

One goal in developing this model is to mitigate the problem of the timescale-dependency of ecological rates. When change is drift-like, as in our simulations (Figs. 2–3), ecological rates are independent of timescale. When change is more bounded, changes over long time intervals will be slower than expected under drift, and fitted drift rates will be correlated with temporal resolution. To assess the degree to which our drift model mitigates the problem of timescale variation in an empirical dataset, we tested the relationship between time interval and rate of change for naïve rates and best-fit drift rates across transitions (that is, contrasts between successive time intervals) in the subset of the Neotoma pollen data chosen above. Because high temporal resolution in the youngest strata coincides with a widely recognized and biologically meaningful increase in ecological rates toward the present (Mottl et al. 2021*a*), we considered only transitions before 2000 years ago, leading to *N* = 70,205 transitions. We quantified naïve rates by calculating the Jaccard dissimilarity across each transition and dividing by the relevant time interval. Both naïve and fitted drift rates were logged, and their relationship with log_10_(time interval) assessed with Spearman rank correlation and linear regression. Data points at the edges of the search window used for maximum-likelihood estimation of drift rate (10^-1^ and 10^-9^; Fig. 6B) were excluded from the regression.

**Fig. 6.**
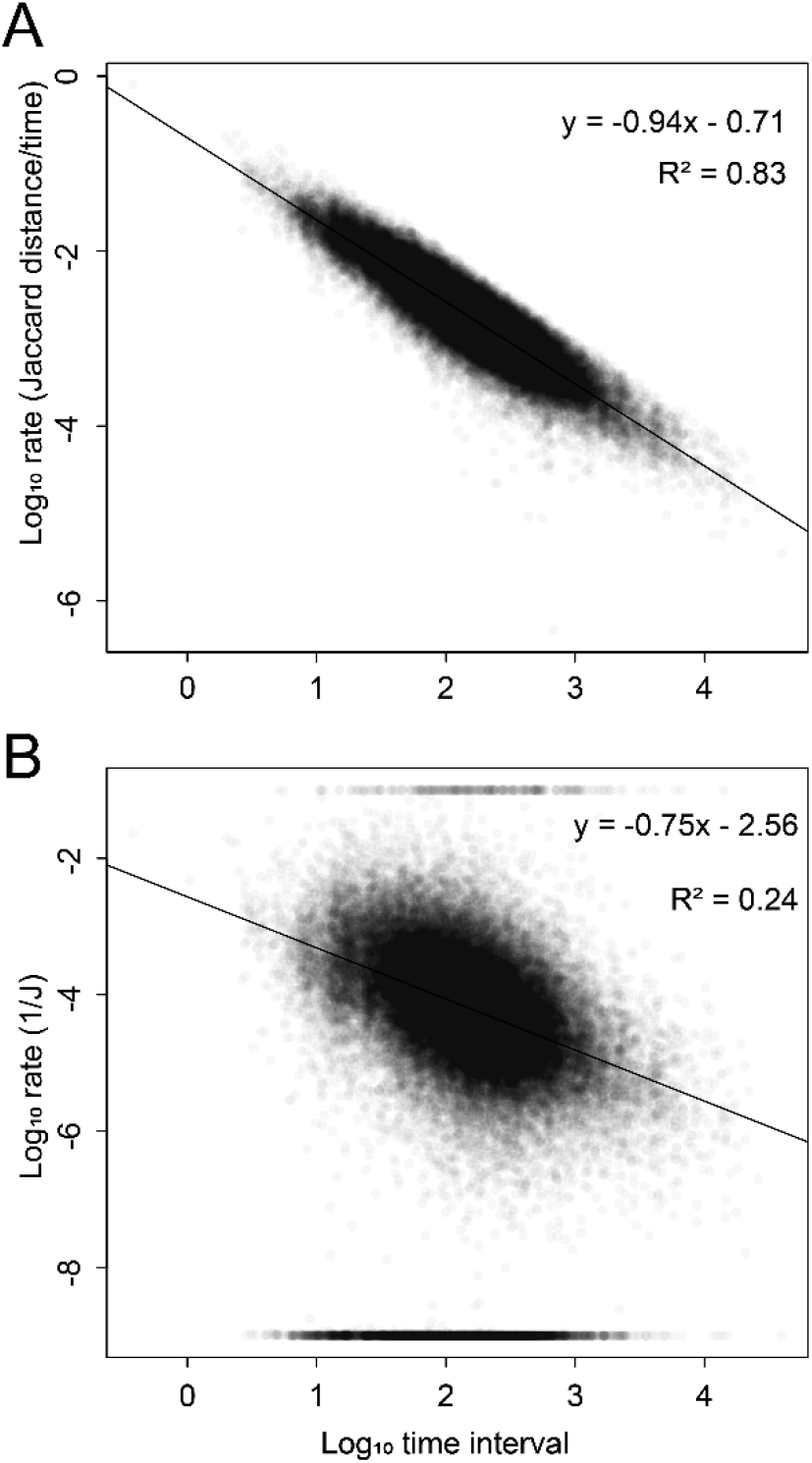
Rate-time scaling of naïve rates and estimated rates of drift across 70,205 transitions in pollen community composition in 1,250 cores from the Neotoma database. **6A**. Jaccard distance over time plotted against time, along with linear regression line. **6B.** Best-fit drift rates plotted against time.

We find naïve rates exhibit a strong negative correlation with timescale, with R^2^ = 0.83 and Spearman’s ρ = 0.90 (Fig. 6A). Nevertheless, the average *amount* of change increases roughly monotonically across the full range of timescales (Fig. S5), and thus it is unlikely the pollen data can be explained by pure community stasis. Relative to naïve rates, fitted 1/*J* values exhibit a weaker relationship with timescale, with R^2^ = 0.24 and ρ = 0.38 (Fig. 6B). By using fitted 1/*J* values to simulate data under a model of drift plus rare migration from a static metacommunity, we find that the residual rate-time scaling in fitted 1/*J* values can be explained by rare migration (Fig. S6). We manually tuned migration rate *m* to find the value that yields a rate-time regression slope closest to the empirical one. We recovered *m* = 0.01 (1% of new individuals sourced from the metacommunity each generation), which is relatively low among published estimates of *m* (Hubbell 2001).

## Discussion

### Why use this model?

We have presented a new approach to measuring and modeling change in communities that is based in biological process: drift resulting from symmetrical competition for limited resources. In using ecological drift, we are not suggesting that it provides an adequate account of how real communities change. Many studies have documented aspects of the natural world that do not comport well with the unified neutral theory of biodiversity (e.g., Ricklefs 2003; McGill et al. 2006; Chisholm and O’Dwyer 2014), suggesting the importance of niche differentiation, predation, competitive asymmetry, and other processes not included in the theory. In lacking migration and speciation, local ecological drift must be even less realistic. Nevertheless, we suggest that this model is uniquely useful for extracting quantitative information about the pace of compositional change, thus facilitating comparisons across taxa, locations, time intervals, and other variables. In this way the model can be useful in similar ways to drift in population genetics, and for similar reasons.

The parameter *J* is directly analogous to the effective population size, *N_e_*, in population genetics: both quantify the stochastic input created by a neutral sampling process (Hu et al. 2006). *N_e_*is not observable in natural populations, but rather is defined by reference to an idealized population with discrete generations, no selection or migration, fixed population size, etc. (Waples 2025). An estimate of *N_e_* from a natural population is not a claim that those idealized conditions are met. Rather, it holds that the potency of genetic drift in the focal population is equivalent to that of an idealized population with the estimated value of *N_e_*. Our interpretation of an estimated *J* is similar: as an “effective community size” (Sloan et al. 2021), it represents the size of an idealized neutral community that would show as much compositional change as the focal community. And, just as the rate of neutral evolutionary change is proportional to 1/*N_e_* (Lande 1976; Hu et al. 2006), 1/*J* is a similarly useful measure of the pace of compositional change (e.g., Figs. 2–6). There are other parallels: for example, faster drift leads to more precise estimates of *N_e_* in population genetics (Waples 2025), and more precise estimates of 1/*J* in our study (Fig. 4).

Applying the drift model to simulated and real communities solves the problem of the rate-time scaling of ecological rates when change is drift-like (Fig. 2), and mitigates the problem even when change is more bounded (Fig. 6). This is because, unlike distance-over-time rate metrics, 1/*J* does not implicitly assume that composition changes linearly with time, which is clearly a less realistic assumption than drift-like changes in relative abundance. In the process of addressing the timescale problem, we have developed solutions to two others. First, empirical sample sizes are often low enough to affect estimates of ecological change (Fig. 2B), especially in paleontological data. Our method allows for unbiased estimation of rates in the face of incomplete sampling (Fig. 2E) because it assumes that observation and true ecological change happen by similar sampling processes. Second, naïve rates are biased upwards in communities with many taxa (Fig. 2C) because more contrasts are possible in more diverse communities. Our approach can accurately estimate true rates in simulated data regardless of taxonomic richness (Fig. 2F) because of the neutrality condition: individuals of different species are ecologically interchangeable in the model, and therefore the predicted abundance changes for a set of individuals do not depend on whether that set of individuals constitutes a single species or many of them. Thus, in addition to being useful for comparing rates between diverse and non-diverse communities, the neutral model is robust to differences in taxonomic resolution. This is especially valuable in paleontological settings where identification to the species level is often impractical, as in pollen data (Figs. 4–5). One final advantage our approach has over many traditional approaches is that it is based on an explicit model of ecological change, and therefore it yields meaningful uncertainties and confidence intervals, returns numbers with a straightforward biological interpretation, and can be flexibly adapted to a variety of hypothesis tests (see Applications).

Perhaps the most important limitation of our implementation of the neutral model is that it ignores abundance transitions that start or end in zero. The likelihood calculation ignores species disappearances because the *J/t* substitution method cannot capture extirpation probability, and it ignores originations because Hubbell’s (2001) speciation parameter does not appear in our model. Importantly, this does not bias rate estimation because our likelihood function is conditioned on not observing those transitions (Fig. S2D). Moreover, transitions to or from zero abundance are likely to be among the least important for studying ecological rates because they commonly represent incomplete sampling rather than true extinctions and originations. Nevertheless, discovering how to better accommodate such transitions would be an important improvement on the model.

There are other approaches to measuring rates of change in community composition outside the two we highlight here. Nogué et al. (2021) ordinated community compositions in each of a set of Holocene pollen sequences from remote islands and regressed ordination scores against time to compare rates of change before and after human arrival. This approach does not produce rates that are comparable between time-series, but sidesteps the timescale problem in the context of a simple hypothesis test. Gibbs et al. (2012) measured rates of change in Paleocene-Eocene nannofossil community composition by smoothing relative abundance trajectories for the most abundant taxa in a time-series, calculating coefficients of variation for each species’s abundance through time across a moving window, and summing coefficients of variation across species. This approach relies on dense sampling and involves lots of user decisions (e.g., how many taxa to retain, the size of the moving window, and details of smoothing), but it successfully captures intervals of high community volatility. Mottl et al. (2021*a*, 2021*b*) dealt with variable temporal resolutions by measuring rates across representative levels randomly chosen from adjacent moving windows of equal duration and repeating many times. This approach is limited by the coarsest resolutions in a time-series and, like the previous example, involves subjective user decisions, but nevertheless performs well in detecting intervals of rapid change in simulated data. Finally, Collins et al. (2000) proposed using the slope of the relationship between log time interval and log ecological dissimilarity (e.g., our Fig. S5) as a measure of the rate of change. While this approach has been adopted in subsequent studies (Korhonen et al. 2010; Pinsky et al. 2025), it arguably confuses change over *time* with change over *timescale*. The slope of the relationship between log interval and log dissimilarity does not measure the rate of change but rather its mode (Gingerich 1993). A low slope indicates bounded change, but bounded change can be rapid (e.g., Fig. 4A); likewise, a high slope indicates directional change, but directional change can be slow (e.g., Fig. 4B).

### Tempo and mode in ecological change

We have proposed a neutral model as a partial solution to the problem of measuring rates of change in community composition. Our solution is based on the supposition that rates of change cannot be meaningfully measured and compared without some model (implicit or explicit) of how change happens: no model, no inference (Sober 1994). In an adjacent field, the development and application of explicit models of change has transformed the study of the tempo and mode of morphological evolution (Uyeda et al. 2011; Hunt et al. 2015; Voje 2016; Hansen 2024). In this field there are three canonical modes of change that can be expressed with just one or two free parameters: directional change (change is proportional to time), stasis (change hovers around an optimum), and drift/random walk (Hunt 2007). Different modes result in different log-log relationships between rates of change and the time interval those rates are measured over: pure directional change results in no relationship between rate and time interval, stasis results in rates inversely proportional to time interval, and random walks plot somewhere in between (Gingerich 1993). These canonical modes of change have analogues in ecology. As in evolution (Voje 2016), sustained directional change in community composition seems to be rare, being limited to especially deterministic succession (Collins et al. 2000); yet, this is the only mode of change under which naïve rates are generally valid. Evolutionary time-series more commonly exhibit patterns ranging from stasis-like to drift-like (Hunt et al. 2025), and ecological time-series may be similar in this regard (Dimichele et al. 2004). Much attention has been paid to long-term ecological stasis (McGill et al. 2005; Reymond et al. 2011), especially in the context of the coordinated stasis (Brett et al. 2025) and turnover-pulse (Vrba 1985) theories. Yet numerous studies report gradual accumulation of ecological change on long timescales (e.g., Overpeck et al. 1992; Bonelli et al. 2006; Cermeño and Falkowski 2009), including this study (Fig. S5), so pure stasis cannot be the whole story. In particular, power spectra of community composition timeseries from a variety of systems are “reddened”: rather than representing white noise around a stable state, change in real communities is most substantial at the lowest frequencies and longest timescales (Pimm and Redfearn 1988; Fastovich et al. 2025).

In our analysis, we find residual rate-time scaling in fitted 1/*J* values (Fig. 6B), indicating that empirical change is more bounded than the expectations of drift. Parallel findings have been reported when fitting drift models to time-series of morphological evolution (Hunt 2012). The scaling pattern in 1/*J* is certainly less severe than that seen in naïve rates, and may be small enough to ignore when variation in time interval is modest or a time-series exhibits a more drift-like pattern of change. In our case studies (Figs. 4–5), variation in time interval does not dictate the primary patterns of inferred rate variation. An investigator seeking to model rate variation as a function of other variables could account for rate-time scaling by including time interval as a predictor of 1/*J*. Alternatively, it may be possible to develop extensions of the drift model that account for the boundedness of real ecological change. Our simulations show that drift with rare migration from a static metacommunity can account for the rate-time scaling in 1/*J* values we observe (Fig. S6). Migration stabilizes communities by rescuing rare species from extinction and drawing species abundances toward their metacommunity values, with metacommunity composition acting like an optimum in an Ornstein-Uhlenbeck model. Migration is not the only possible cause of community stasis, and indeed niche stabilization is frequently posited as a cause of stasis via negative frequency dependence in species abundances (McGill et al. 2005; Adler et al. 2007; Reymond et al. 2011). Nevertheless, developing a full likelihood model of drift with migration might allow for the estimation of ecological rates while solving the timescale problem altogether. Such a model could even be used phenomenologically to test possible causes of community stasis, such as whether ecological stasis was more prominent in more stable abiotic settings.

### Applications

We have presented a method for measuring and modeling change in community composition that is based on the process of ecological drift, produces numbers that have biological meaning and are comparable across datasets, and is relatively robust to variation in timescale, sampling, and taxonomic resolution (Fig. 2). Models based on drift have become central to the study of morphological evolution in phylogenetics (Huey et al. 2019) and paleobiology (Goswami and Clavel 2025), and we suggest they could achieve similar results in paleoecology. The recent widespread availability of abundance databases like BioDeepTime (Smith et al. 2023) makes the development of models for making robust inferences from ecological time-series especially timely. We conclude by highlighting five promising applications for drift models in deep-time community ecology:

1. *Descriptive/exploratory paleoecology.* By mitigating non-biological biases in the measurement of ecological rates, our method can be useful for exploring and testing the significance of apparent patterns of ecological change, e.g., through comparing rates across sites (Fig. 4) or testing for rate shifts (Fig. 5).
2. *Testing covariates of ecological rates.* Fig. 5 shows how fitted drift rates can be used to investigate the timing of increases in ecological rates associated with human impacts. Future work could more explicitly build and test models in which drift rates vary as a function of possible drivers of change. For example, pollen assemblage data could be combined with measures of climate, land use, human population density, and other factors to understand the proximate causes mediating anthropogenic impacts on ecological communities.
3. *Coordinated stasis.* Models of drift and related processes have brought precision and analytical rigor to the debate over punctuated equilibrium (Hunt et al. 2025), and drift models could contribute to the theory of coordinated stasis in a similar way. For example, Strotz and Lieberman (2021) tested a strict form of coordinated stasis by asking whether a set of observed changes in brachiopod community composition could be explained purely by sampling noise. This is essentially a special case of our model in which *J* is infinite, and in our framework such a model could be contrasted against a single-rate model, or a model in which stretches of little or no change are punctuated by rapid change, or other possible alternatives. Our method could also provide quantitative tests of the prediction from coordinated stasis that intervals of ecological upheaval coincide with rapid morphological and taxonomic change (Brett et al. 2025).
4. *Tests of neutral theory.* This study uses neutral theory because it provides a convenient rate metric, not because it is true. However, neutral models do yield parsimonious explanations for a broad range of natural phenomena (Rosindell et al. 2012; Saulsbury et al. 2024), and our model could be used to test the importance of drift and dispersal limitation in real ecosystems. For example, an investigator could test whether values of parameters in the theory (e.g., community size, migration rate) estimated from community time series agree with estimates from species abundance distributions. Additionally, because neutral theory predicts that drift should be stronger in smaller communities, one could test whether estimated drift rates from time-series correspond to proxies of community size, such as pollen accumulation rate.
5. *Macroecology.* Much effort in macroecology has gone into understanding how and why biological properties such as body size (Brown 1995), range size (Tomašových et al. 2015), and diversification rate (Rabosky et al. 2018) vary across major spatial and environmental gradients around the globe. It remains relatively unexplored how the tempo and mode of ecological change vary across these gradients. Is ecological change faster on land or in the sea? In shallow or deep marine settings? In or outside the tropics? Do patterns of ecological change differ across major divisions in the tree of life (Montgomery et al. 2026), e.g. between vertebrates and phytoplankton? Do ecological rates correspond with evolutionary rates, e.g., rates of speciation or extinction? Answering these and related questions will depend on our ability to reliably measure and model ecological change across heterogeneous datasets. In this the application of simple, extensible process models based in sound ecological theory is a promising way forward.

