## Supplementary material for "Measuring and modeling ecological rates with neutral theory": SI

**Supplementary Information for: Measuring and modeling ecological rates with neutral theory**

**Combining change in multiple species using conditional probability**

Step 3 of the implementation of our model (main text) involves expressing a multinomial distribution as the joint probability of multiple conditional binomial distributions. Here we show what that looks like in more detail. The probability that a draw of size *n* with replacement from *S* categories with probabilities $p\bar{p}$ results in a set of successes within each category in $k\bar{k}$ is given by the multinomial distribution $M(n,p)$:

$$M(n,\bar{p},\bar{k})=P\left( \bar{k}|n,\bar{p} \right)$$

$$Pr\left( k|n,p \right)=Pr(X_{1}=k_{1} and\ldots and X_{S}=k_{S})$$

$$X\sim M(n,p)$$

The joint probability above can be decomposed into the product of conditional probabilities:

$$P\left( \bar{k}|n,\bar{p} \right)=P\left( k_{1} \right|n,p_{1})\cdot P\left( k_{2} | n,p_{1\to2},k_{1} \right)\cdot\ldots\cdot P\left( k_{j} | n,p_{1\to l},k_{1\to j-1} \right)$$

$$Pr\left( k|n,p \right)=\prod_{i=1}^{S} Pr\left( k_{i}|n,p_{1\to i},k_{1\to i-1} \right)$$

The probability of a draw in category *i* conditional on the draws in categories $1\to i-1$ is the same as the probability of a binomial draw in category *i* with categories $1\to i-1$ removed from consideration. The above probability can be expressed as a product of probabilities under the binomial distribution as follows:

$$P\left( \bar{k}|n,\bar{p} \right)=B\left( n,p_{1},k_{1} \right)\cdot B\left( n-k_{1},\frac{p_{2}}{1-p_{1}},k_{2} \right)\cdot\ldots\cdot B\left( n-\sum_{i=1}^{j-1} k_{i},\frac{p_{j}}{1-\sum_{i=1}^{j-1} p_{i}},k_{j} \right)$$

$$Pr\left( k|n,p \right)=\prod_{i=1}^{S} Pr(X_{i}=k_{i})$$

$$X_{i}\sim B\left( n-\sum_{j=1}^{i-1} k_{j},\frac{p_{i}}{1-\sum_{j=1}^{i-1} p_{j}} \right)$$

Eqn. 3 in the main text has the same form as the above equation.

**Combining sampling and true change into a single step**

Consider a transition in composition in a neutral community with size $J=j_{1}$ over a single generation. Suppose also that the community has been observed by sampling with replacement before and after this transition, with sample size $j_{2}$ before the transition and sample size $j_{3}$ after. As justified in the main text, we incorporate these sampling steps by treating them as if they were additional phases of drift. Using the *J/t* substitution described in the main text, these three phases of drift with sizes $j_{1}$, $j_{2}$, and $j_{3}$ can be substituted with drift in a community of size $j_{1}j_{2}j_{3}$ over $j_{2}j_{3}$, $j_{1}j_{3}$, and $j_{1}j_{2}$ timesteps, respectively. These three phases combine into a single phase with

$$\frac{J}{t}=\frac{j_{1}j_{2}j_{3}}{j_{2}j_{3}+j_{1}j_{3}+j_{1}j_{2}}=\frac{j_{1}j_{2}j_{3}}{\frac{j_{1}j_{2}j_{3}}{j_{1}}+\frac{j_{1}j_{2}j_{3}}{j_{2}}+\frac{j_{1}j_{2}j_{3}}{j_{3}}}=\frac{1}{\frac{1}{j_{1}}+\frac{1}{j_{2}}+\frac{1}{j_{3}}}$$

This last expression appears in main text Equation 3.

**Supplementary Figures**

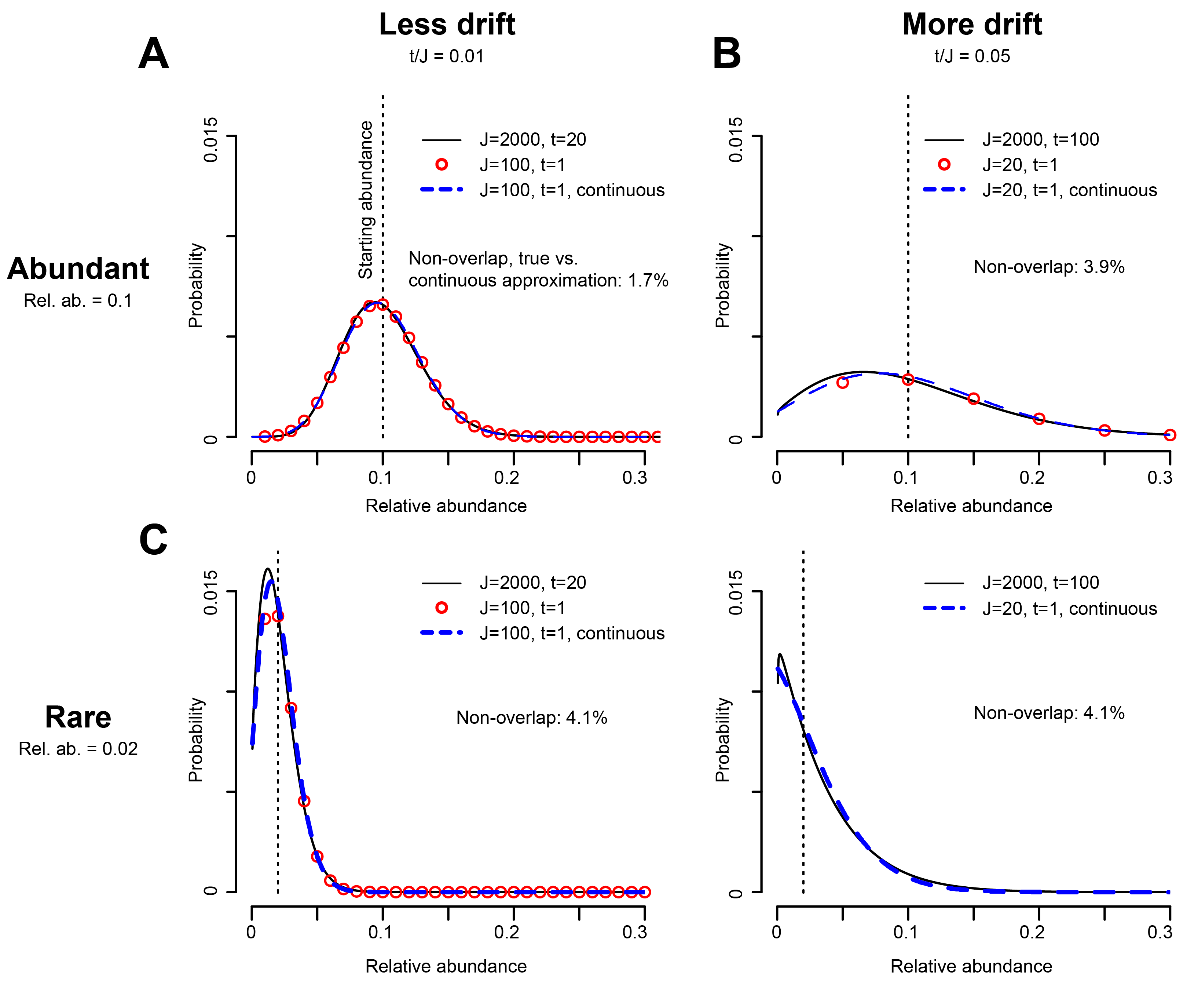

**Fig. S1.** Accuracy of the *J/t* substitution method for estimating the probability of abundance changes under neutral theory. Plotted after main text Fig. 1A. Error quantified as the percent non-overlap between the true (black) and estimated (blue dashed) probability distributions. Probabilities from the *J/t* approximation have been multiplied by a factor for comparison with true distribution (see main text Fig. 1A). All probabilities are conditional on not observing extinction. The *J/t* approximation becomes somewhat less accurate under lower starting abundances and higher drift, although the relationship is not linear. **S1A.** Low drift and high relative abundance. **S1B.** High drift, high relative abundance. **S1C.** Low drift and low relative abundance. **S1D.** High drift and low relative abundance. Note that the approximation underestimates the probability of extinction here, but overestimates it in S1B and S1C. Discrete binomial approximation not shown here because relative abundance 0.02 is not defined for a community of size *J*=20.

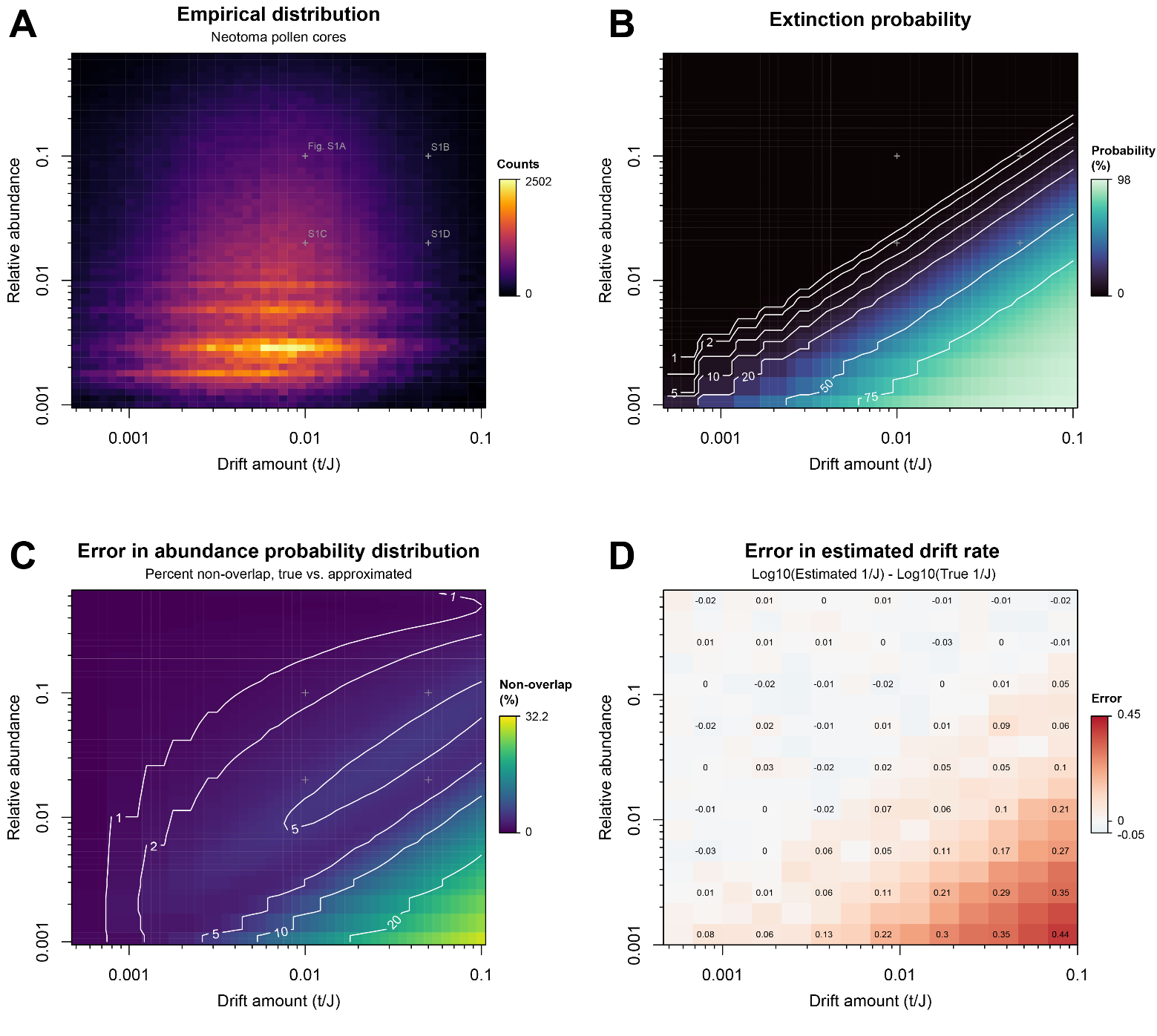

**Fig. S2.** Method performance across an empirically determined range of ecological rates and relative abundances. **S2A.** Relative abundances and estimated ecological drift amounts (*t/J*) from Neotoma pollen cores. Drift amounts obtained by fitting 1/*J* separately to every change in community composition in the dataset and multiplying by the relevant time interval in years; plot encompasses the 4.9^th^ to 98.2^th^ percentiles. Relative abundances encompass the 4.4^th^ to 99.3^th^ percentiles of all 1.8 million non-zero relative abundances in the dataset. Grey points indicate positions in parameter space of cases presented in Fig. S1. **S2B.** True extinction probability as a function of drift amount and relative abundance, as determined using a transition matrix of size *J*=2000. Extinction is most probable at low abundances and under high drift. **S2C.** Error in estimating abundance probability distributions, as quantified by percent non-overlap between true and estimated probability distributions (c.f. Fig. S1). True probabilities again determined using a transition matrix of size *J*=2000. Non-overlap exceeds 10% when extinction probability exceeds ~50%, but non-overlap never exceeds 20%. **S2D.** Error in estimating drift rate across parameter space, as quantified by difference between estimated and true drift rate in log_10_ space. For each cell, a two-species community of size *J*=20,000 was simulated for *t* timesteps and only the first and last timesteps were retained; this was repeated 2000 times and *fitJ*() was run on this set of simulations. The true drift rate is within the 95% confidence interval in most cases except for the bottom right corner of parameter space. Drift rate is overestimated when extinction probability is high because the approximated likelihood tends to underestimate the probability of extinction in this part of parameter space (Fig. S1D).

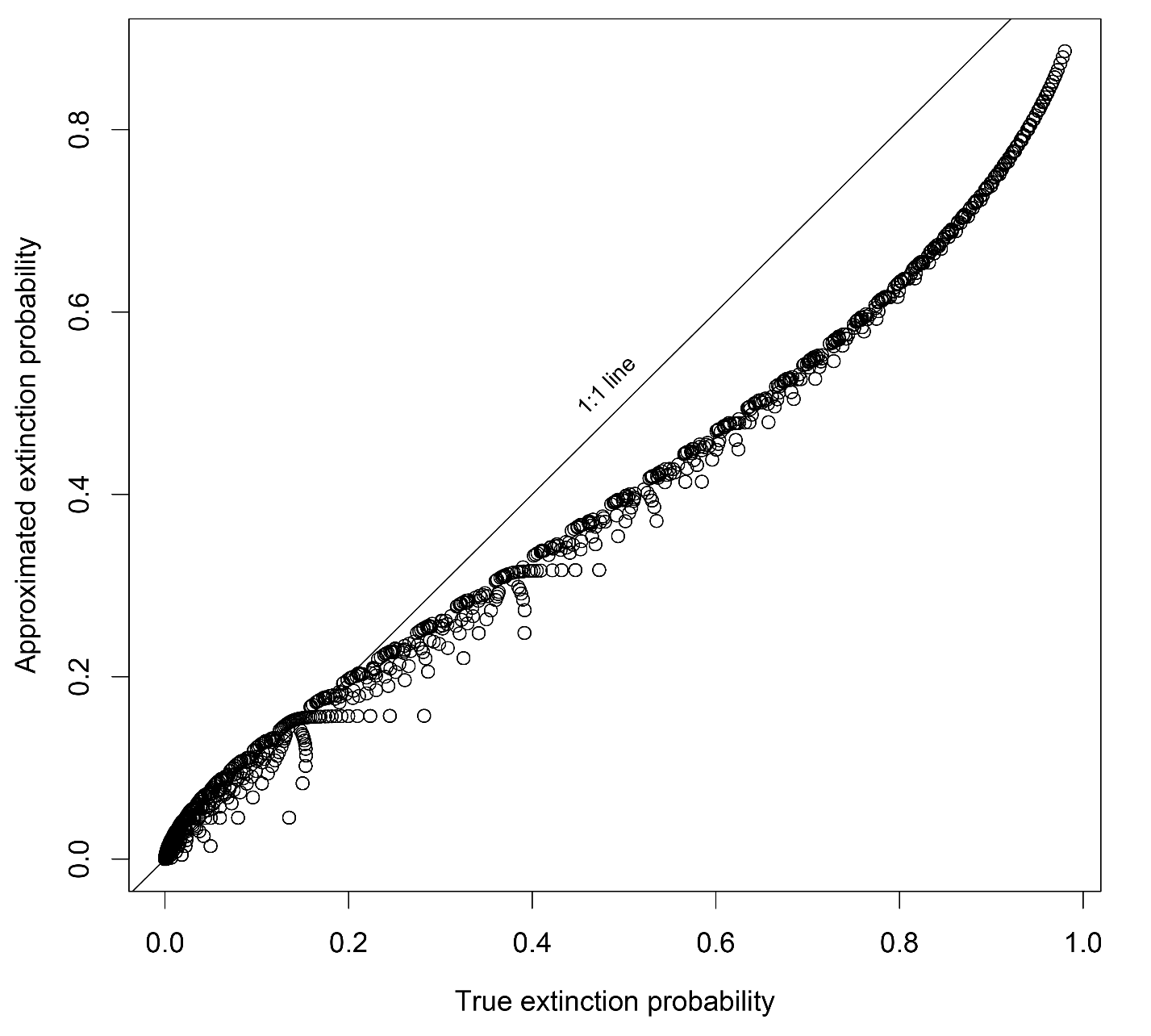

**Fig. S3.** True extinction probability versus one minus the denominator in main text Fig. 2, which closely approximates extinction probability. Regressing approximated versus true extinction probability yields y = 0.803x + 0.00215, R^2^ = 0.993.

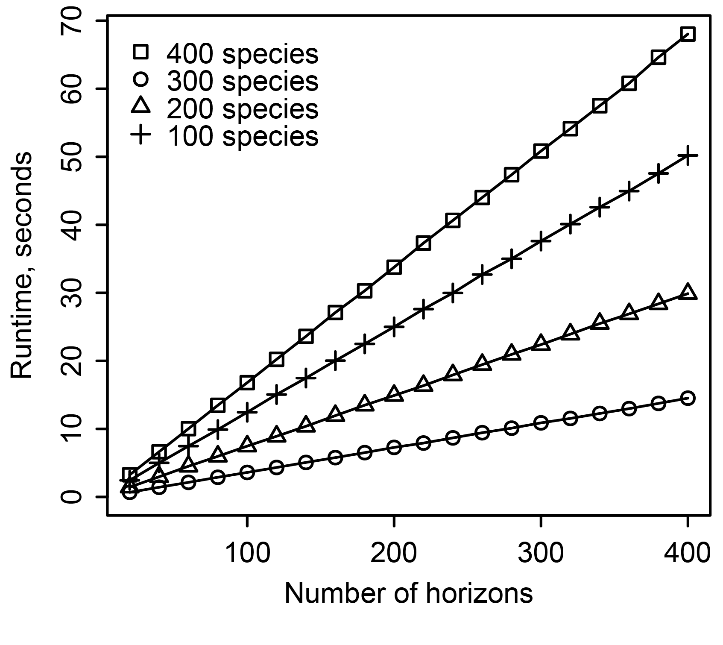

**Fig. S4.** Runtime of the fitJ() function.

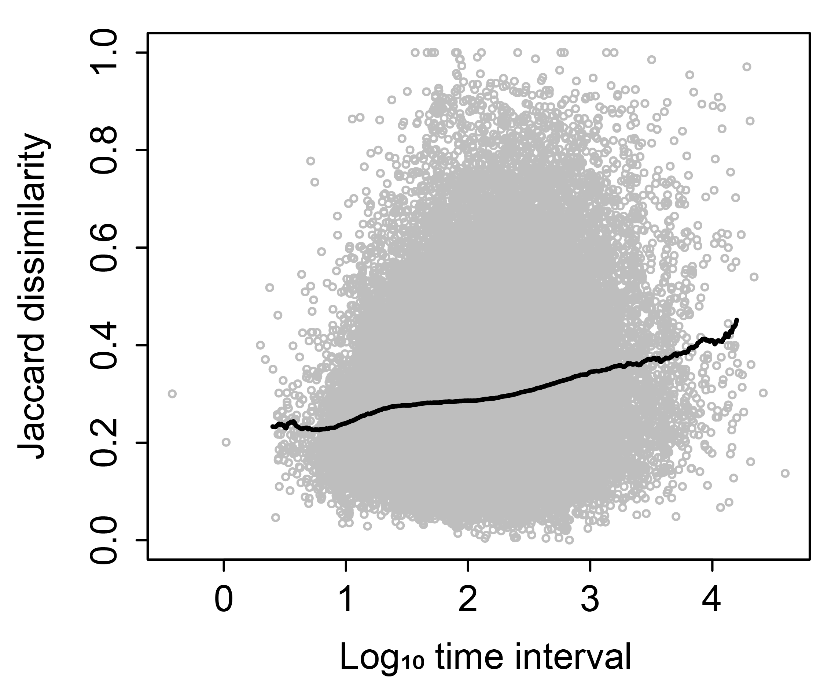

**Fig. S5.** Jaccard dissimilarity plotted against time interval for 70,205 transitions in pollen community composition from the Neotoma dataset (grey), along with mean dissimilarity calculated in a sliding window of width 0.2 in log space (black).

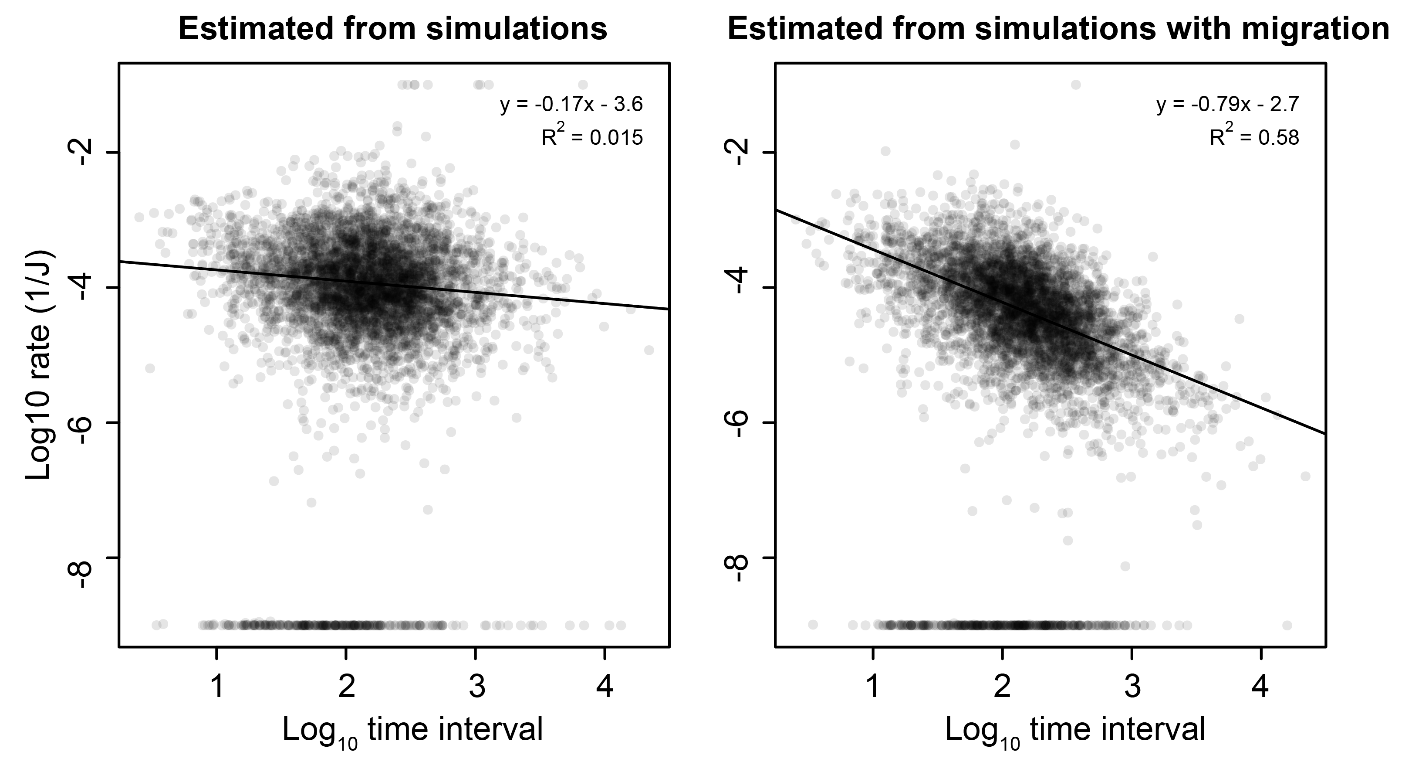

**Fig. S6.** Migration as a possible cause of rate-time scaling in estimated drift rates. We randomly sampled 5,000 community transitions with associated fitted 1/*J* values from the Neotoma dataset and permuted the associated time interval values to remove rate-time scaling. We used 1/*J* values and permuted time intervals to simulate new data with or without migration, then fit J to simulated data. We show the migration rate that resulted in a regression slope most similar to that in main text Fig. 6B. Linear regressions ignore zero-rate data points due to their much greater abundance here than in main text Fig. 6 (~10% vs. ~1%). Left, drift rates fitted to data simulated without migration. Right, drift rates simulated under the effect of rare (m = 0.01) migration from a metacommunity with relative abundances equal to those of the local community at the start of the simulation.
